# Design of SARS-CoV-2 RBD mRNA Vaccine Using Novel Ionizable Lipids

**DOI:** 10.1101/2020.10.15.341537

**Authors:** Uri Elia, Srinivas Ramishetti, Niels Dammes, Erez Bar-Haim, Gonna Somu Naidu, Efi Makdasi, Ofer Cohen, Dan Peer

## Abstract

The novel coronavirus SARS-CoV-2 has been identified as the causal agent of COVID-19 and stands at the center of the current global human pandemic, with death toll exceeding one million. The urgent need for a vaccine has led to the development of various immunization approaches. mRNA vaccines represent a cell-free, simple and rapid platform for immunization, and therefore have been employed in recent studies towards the development of a SARS-CoV-2 vaccine. In this study, we present the design of a lipid nanoparticles (LNP)-encapsulated receptor binding domain (RBD) mRNA vaccine. Several ionizable lipids have been evaluated *in vivo* in a luciferase mRNA reporter assay, and two leading LNPs formulation have been chosen for the subsequent RBD mRNA vaccine experiment. Intramuscular administration of LNP RBD mRNA elicited robust humoral response, high level of neutralizing antibodies and a Th1-biased cellular response in BALB/c mice. These novel lipids open new avenues for mRNA vaccines in general and for a COVID19 vaccine in particular.

## Introduction

Severe acute respiratory syndrome coronavirus (SARS-CoV) 2, is a novel coronavirus identified as the etiological agent of coronavirus disease 19 (COVID-19). This coronavirus stands at the center of the current global human pandemic, with recent reports of more than 36 million cases and over one million deaths worldwide [1]. The urgent need for a vaccine has led to an unprecedented recruitment of academic laboratories, hospitals and pharmaceutical companies around the world, which translated into a wide array (>180) of pre-clinical and clinical studies being conducted in an effort to develop an effective vaccine against SARS-CoV-2 [2]. These vaccine candidates can be classified into several categories: inactivated/live attenuated virus, recombinant viral vector, recombinant protein, DNA vaccine and messenger RNA (mRNA) vaccine.

The mRNA vaccine platform has developed extremely rapidly in the past few years, mainly due to advances in mRNA stabilization and the introduction of efficient delivery methods that originated largely from the siRNA field. mRNA vaccines hold several advantages over traditional vaccine approaches such as inactivated/ live attenuated, subunit or DNA-based vaccines: This platform poses no potential risk of infection or genome integration, does not require entry to the nucleus, and can be developed very rapidly and easily. This last advantage has been demonstrated very clearly in the current COVID-19 pandemic, with the development of an mRNA vaccine by Moderna directed against the spike protein of SARS-CoV-2, with only 66 days from sequence selection to first human dosing [3].

One of the main challenges in mRNA therapy is efficient delivery of mRNA to target cells and tissues. High susceptibility to degradation by omnipresent ribonucleases (RNases), together with inherent negative charge, hinder the successful delivery of mRNA to cells and subsequent translocation across the negatively charged cell membrane. Hence, successful mRNA delivery requires a carrier molecule which will protect it from degradation, and facilitate cellular uptake. Lipid nanoparticles (LNPs) are a clinically advanced, non-viral delivery system for siRNA, approved by the FDA [4]. In the past few years, LNPs have emerged as one of the most advanced and efficient mRNA delivery platforms. Recent reports demonstrate antigen-encoded mRNA encapsulated in lipid nanoparticles (mRNA-LNPs) as a potent vaccine platform for several infectious diseases including viral infections such as HIV, CMV, Rabies, influenza, Zika, and most recently SARS-CoV-2 [3,5–10]. LNPs are generally comprised of four components: 1) an ionizable lipid, which promotes self-assembly of the LNPs; 2) cholesterol as a stabilizing agent; 3) a phospholipid for support of lipid bilayer structure; and 4) a polyethylene glycol (PEG)-lipid, which increases the half-life of the molecule. Of these, the ionizable lipid is a key player for efficient intracellular delivery of the mRNA. The ionizable lipid facilitates mRNA encapsulation, promotes interaction with cell membrane, and is suggested to play a part in endosomal escape, a key step in the delivery of mRNA into the cytoplasm, and subsequent translation of the protein of interest [11].

Our lab previously demonstrated the design and synthesis of novel ionizable amino lipids for efficient siRNA delivery to leukocyte subsets [12]. Herein, we have selected several structurally different ionizable lipids and screened for *in vivo* mRNA-LNPs delivery for vaccine applications. The screen yielded two LNP formulations, which were chosen for further immunization studies using SARS-CoV-2 RBD mRNA. These experiments demonstrated the development of a specific humoral and cellular response against the antigen, as well as neutralizing antibodies that blocked viral infection in a VSV Plaque Reduction Neutralization Test (PRNT). Additionally, we measured Th1/Th2 specific cytokine secretion in response to LNP-RBD mRNA vaccination.

## Results

### LNPs preparation and physicochemical characterization

The structures of the new amino lipids are shown in Figure 1A. The lipids were structurally different in their head group region, linker and lipid tails. LNPs were produced by mixing of lipids and mRNA through micro fluidic mixture device. A schematic illustration of LNP synthesis is shown in Figure 1B and described in details in the materials and methods section. The resultant mRNA-LNPs were small and uniformly distributed as evidenced by small hydrodynamic diameter and polydispersity index (PDI), measured by dynamic light scattering (DLS) (Figure 1C). Except lipid 5, the mean size of the LNPs was less than 100 nm in diameter. Transmission electron microscopy (TEM) analysis supported the DLS data, showing small and uniform size distribution of the particles (Figure 1D).

**Figure 1.**
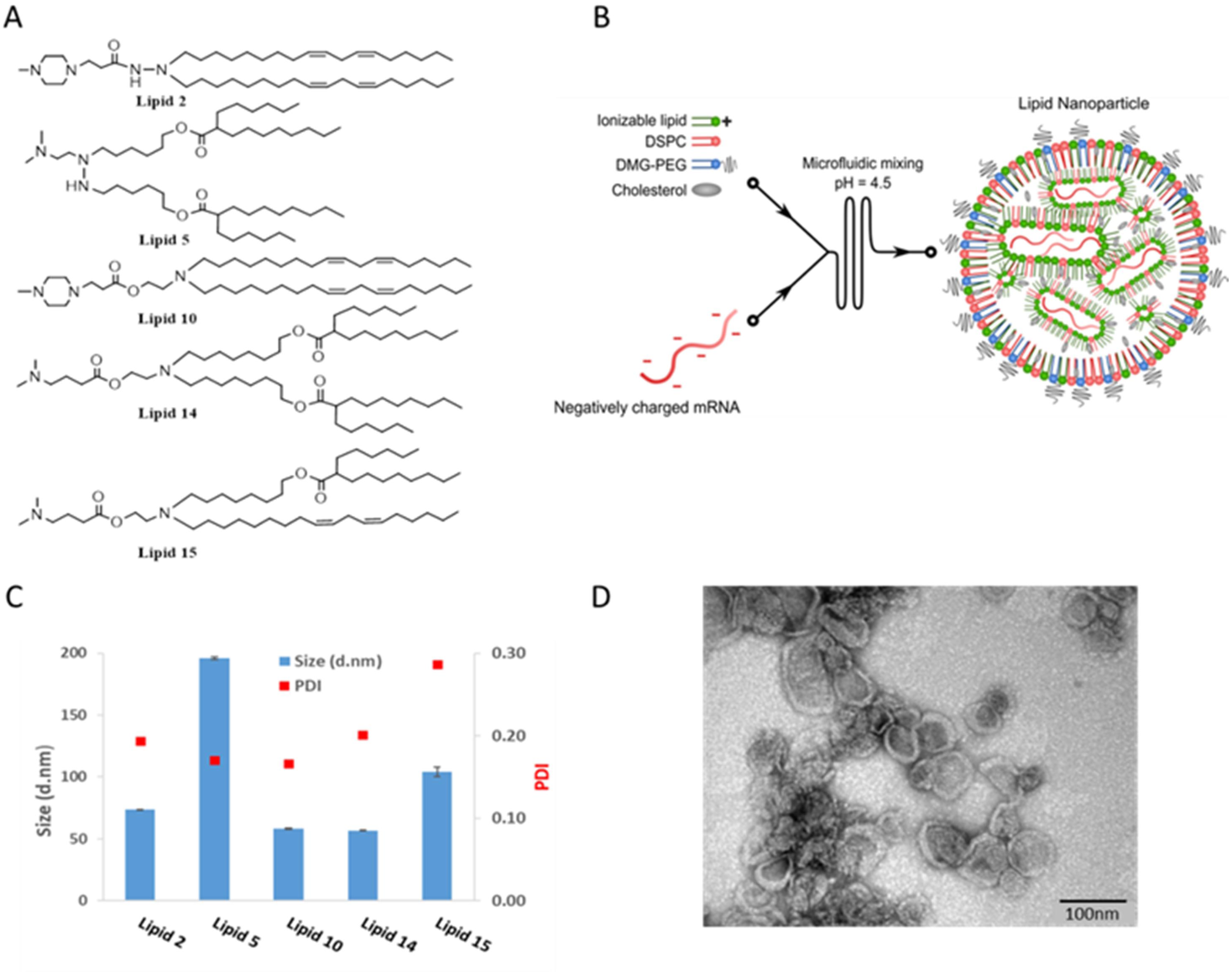
Chemical structures and physicochemical properties of designed LNPs. Panel A: Schematic illustrations of the structures of the designed lipids. Panel B: Schematic illustration of LNP synthesis. Panel C: Representative size distribution and polydispersity index (PDI) of LNPs measured by dynamic light scattering Panel D: Representative TEM images of LNP #2. Scale bar 100 nm.

### *In vivo* luciferase expression screen shows two distinct formulations with superior protein expression

In order to evaluate the *in vivo* efficiency of the LNPs in terms of distribution, protein expression efficiency and kinetics, we conducted a luciferase mRNA-based *in vivo* screen. Animals were injected via the intradermal (i.d.), intramuscular (i.m.) or subcutaneous (s.c.) routes with 5ug luciferase-mRNA encapsulated with one of the five LNP-based formulations, represented herein as LNP #2, #5, #10, #14, #15, and luciferase expression was evaluated daily using IVIS. As shown in Figure 2, LNPs #14 and #15 were superior to other formulations in terms of protein expression level and its duration in all three routes of administration. Since i.d. and i.m. injections exhibited higher and more prolonged protein expression, further immunization studies were conducted using these routes of administration. We chose to proceed with LNPs #2, #14, and #15. Although not showing relative advantage in terms of protein expression *in vivo*, we chose to include LNP #2 in order to eliminate the possibility of a discrepancy between *in vivo* luciferase expression and the resulting immunologic response.

**Figure 2.**
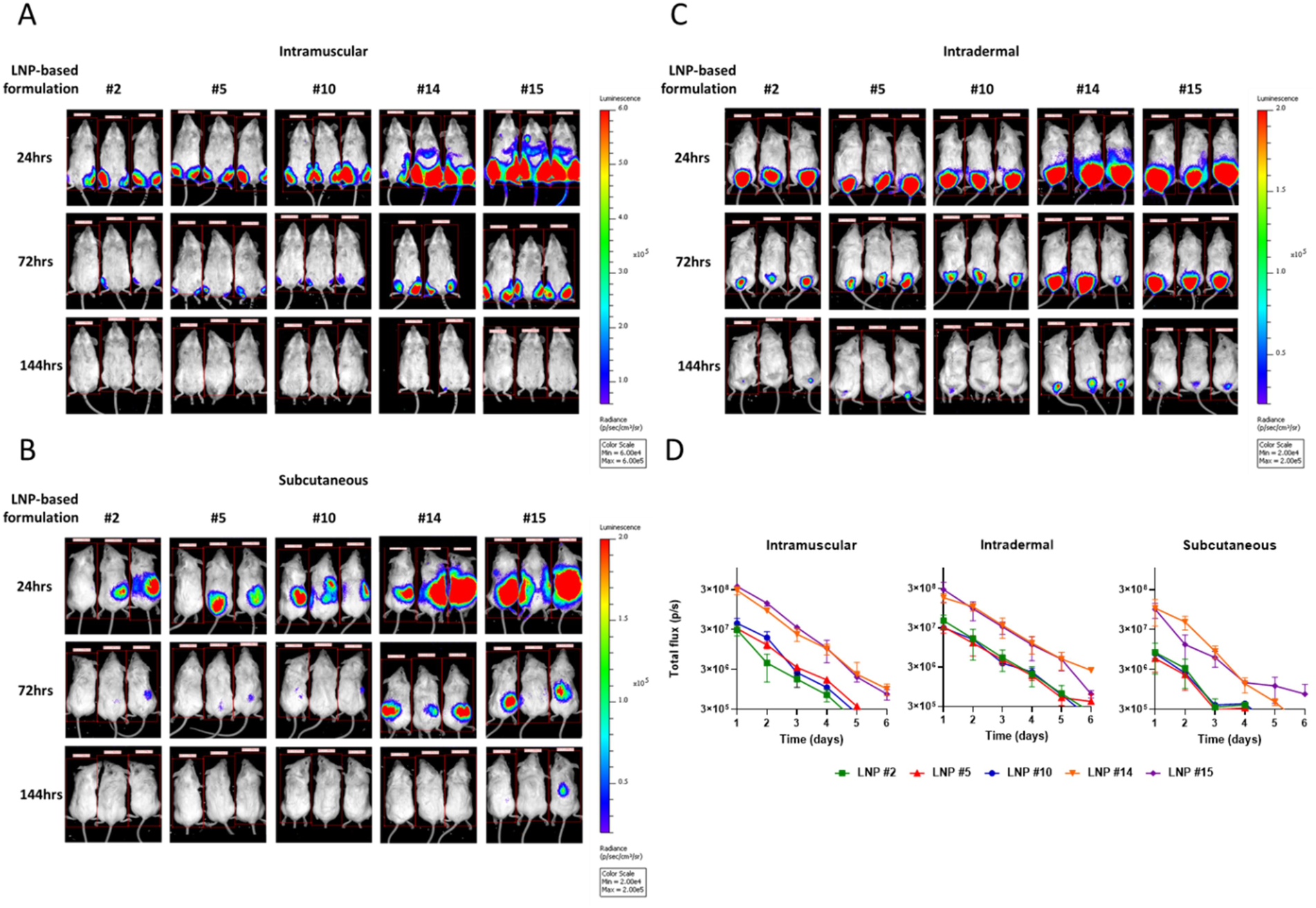
*In vivo* expression pattern of LNP-encapsulated luc mRNA. Representative IVIS images of groups of female BALB/c mice injected with 5 μg luc mRNA encapsulated by five LNP formulations by the intramuscular (panel A) intradermal (panel B), and subcutaneous (panel C) routes. Panel D-quantification of the bioluminescent signal detected throughout six days of monitoring.

### Immune response to luciferase expression

Next, we examined the immune response that was developed against the luciferase protein. Intramuscular-immunized mice were bled and sacrificed 4 weeks after immunization with LNP-encapsulated luciferase mRNA. A panel of assays including ELISA for the detection of anti-luciferase antibodies, and ELISpot for evaluation of luciferase-specific cellular response were tested. While the humoral response was limited, with no statistically significant differences between the vaccinated animals and the control naïve mice, as expected after a single dose administration (Figure 3A), a substantial cellular response was detected when splenocytes were stimulated with the luciferase protein. Interestingly, immunization with LNPs #2 and #14 yielded a significantly stronger (∼3 fold and ∼5 fold, respectively) cellular response compared to LNP #15 (Figure 3B). Based on these results, we decided to perform the following vaccinations with the two leading formulations #2 and #14.

**Figure 3.**
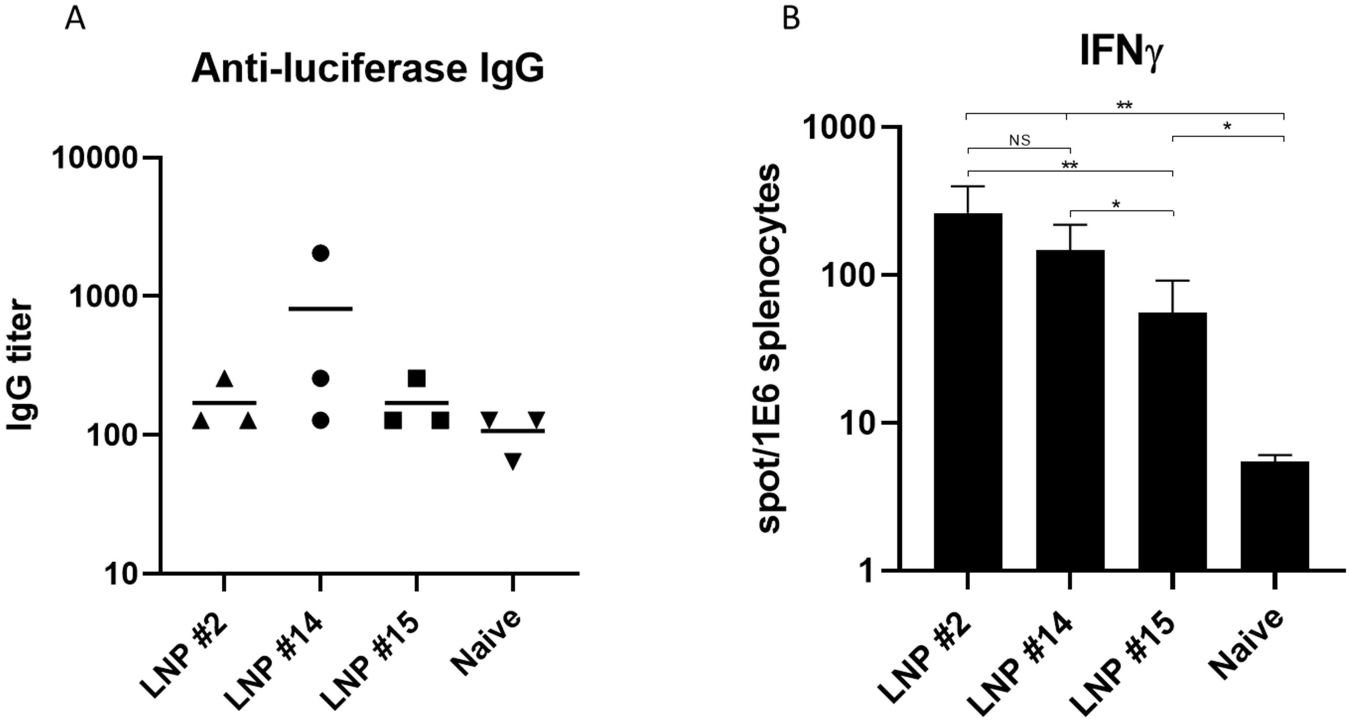
*In vivo* luciferase expression leads to specific cellular immune response. Female BALB/c mice were intramuscularly administered with 5μg LNPs-luc mRNA or untreated (naïve). Serum and spleens were collected 28 days post-administration for evaluation of luciferase-specific humoral (panel A) and cellular response (panel B), as described in the materials and methods section. Statistical analysis was performed using unpaired two-tailed Student’s t-test (n.s., not significant; *p<0.05, **p<0.01).

### Immune responses in RBD mRNA-vaccinated mice

Similar to SARS-CoV, SARS-CoV-2 recognizes angiotensin-converting enzyme 2 (ACE2) as receptor for host cell entry. SARS-CoV-2 spike (S) protein consists of S1, including receptor-binding domain (RBD), and S2 subunits [13]. For our vaccine platform, we chose the RBD of SARS-CoV-2 as the target antigen for the mRNA coding sequence, as described in the materials and methods section. Mice were immunized i.m. or i.d. with either naked RBD mRNA (5μg), LNPs-encapsulated (#2 or #14) RBD mRNA (5μg), or Empty LNPs. We also included a recombinant RBD (rRBD) study group, which was immunized (s.c.) with a recombinant RBD protein (10μg) for comparison with the mRNA vaccinated groups. In all groups, a prime-boost vaccination regimen was employed, with animals being primed at day 0, and boosted 25 days later. Blood and spleens were collected at day 23 (pre-boost) and 39 (14 days post-boost) for evaluation of immune responses (see outline in Figure 4A).

**Figure 4.**
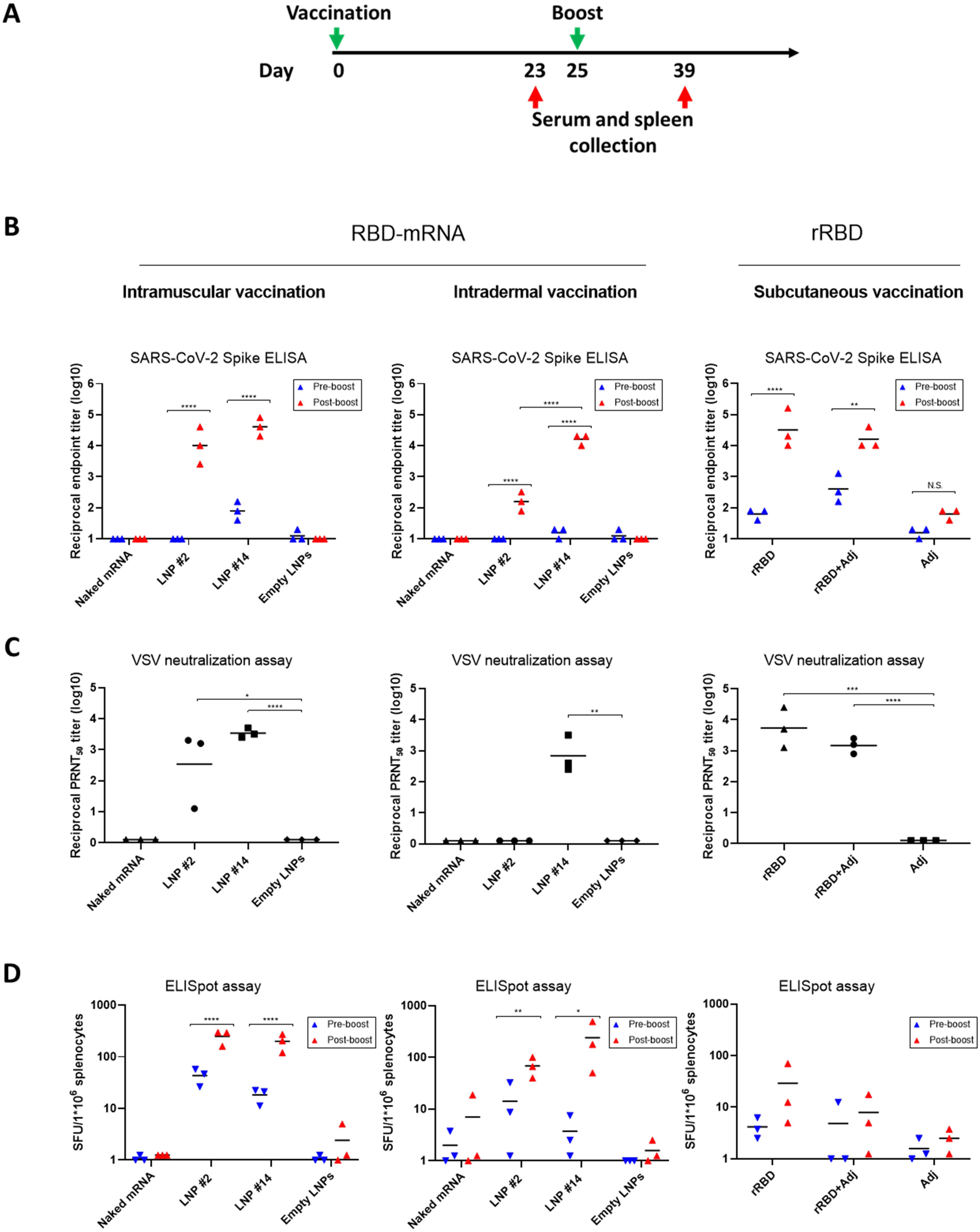
Immunization of mice with LNPs-RBD mRNA leads to a robust immune response. Female BALB/c mice were immunized either i.m. or i.d. with 5μg LNPs-RBD mRNA or s.c. with 10μg rRBD, and boosted with an equivalent dose 25 days later. Serum and spleen were collected at days 23 (“pre-boost”) and 39 (“post-boost”) after initial vaccination. Panel A-Schematic diagram of immunization and sample collection. Panel B-SARS-CoV-2 spike-specific IgG antibody titer was determined by ELISA. Panel C-PRNT_50_ titers were determined post-boost using a VSV-based pseudovirus PRNT assay. Panel D-SARS-CoV-2 spike-specific cellular response was determined by ELISpot. Statistical analysis was performed using a two-way ANOVA with Tukey’s multiple comparisons test (for ELISA data) or an unpaired two-tailed Student’s t-test (for PRNT_50_ and ELISpot data) (*p<0.05, **p<0.01, ***p<0.001, ****p<0.0001).

As can be seen in Figure 4B, pre-boost humoral response against SARS-CoV-2 spike was limited, both in mRNA (naked or LNP-encapsulated) and rRBD-vaccinated mice. However, a robust antibody response could be detected 14 days after the boost in both LNP-encapsulated mRNA and recombinant protein groups, while no response was observed in the naked mRNA group. While both LNP formulations exhibited a boost effect at the i.m. route, a differential antibody response was observed at the i.d. route between LNP #2-and #14-encapsulated RBD mRNA, with only the latter yielding a substantial anti-RBD titer (>10,000) (Figure 4B). Most importantly, this differential response was also evident in the post-boost VSV neutralizing assay, where i.d. LNP #14-encapsulated RBD mRNA vaccination induced a significant neutralizing response, while no neutralizing activity was apparent in LNP #2-encapsulated RBD mRNA vaccinated mice (Figure 4C). Immunization with rRBD also led to a robust boost effect in terms of anti-spike and neutralizing antibodies. Interestingly, no significant difference was observed between vaccinations with the recombinant RBD protein alone or in the presence of the CFA/IFA adjuvant.

The cellular immune response to SARS-CoV-2 plays a crucial role in the ability of the immune system to overcome infection [14, 15]. We thus evaluated the cellular response that developed after immunization with LNP-encapsulated RBD or rRBD by using the ELISpot method for quantification of IFNγ-secreting cells. In contrast to the humoral response, which was very limited before boost administration, a clear specific cellular response was observed 23 days after priming, particularly in mice that were vaccinated via the i.m. route.

A significant increase in specific cellular response was observed after boost administration in both i.m. and i.d. routes of administration, and in both LNP formulations of the RBD mRNA. Conversely, immunization of mice with recombinant RBD did not lead to a significant cellular response, and the post-boost elevation in IFNγ secretion was not statistically significant (Figure 4D).

We next evaluated the Th1/Th2 cytokine secretion profile of LNP RBD mRNA vaccinated mice. As shown in Figure 5 (and Figure 4D of ELISpot results for IFNγ response), a specific and statistically significant secretion of IFNγ and IL-2 was observed in vaccinated mice compared to vehicle treatment, both before and after boost administration. In contrast, Th2 cytokines were either below limit of detection (IL-4) or in comparable levels in vaccinated versus vehicle-treated animals (IL-10).

**Figure 5.**
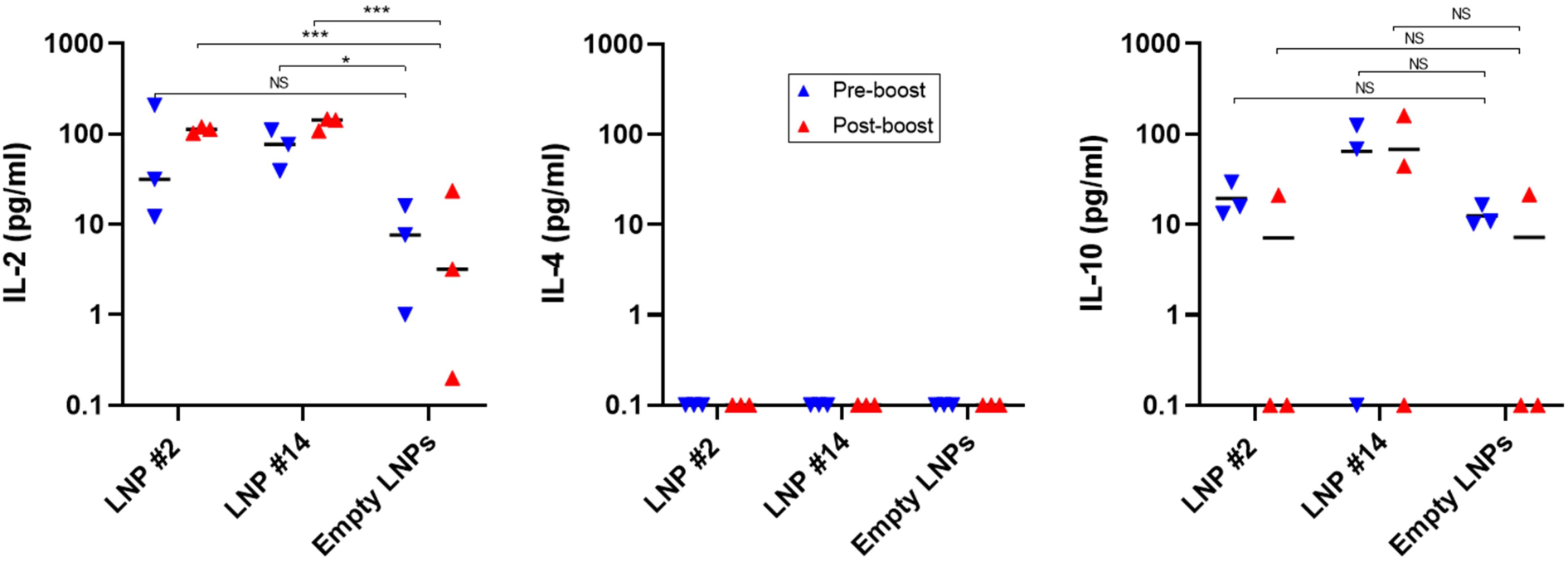
Cytokine profile of induced responses. Splenocytes from i.m. vaccinated mice were stimulated with SARS-CoV-2 spike and analyzed for cytokine secretion by ELISA. Statistical analysis was performed using an unpaired two-tailed Student’s t-test (*p<0.05, **p<0.01, ***p<0.001, ****p<0.0001).

## Discussion

Severe acute respiratory syndrome coronavirus 2 (SARS-CoV-2) has recently emerged as a global pandemic, risking most of the earth’s population health, and leading towards a worldwide economic crisis since its outbreak. Numerous vaccination platforms have been recently employed in the quest for an effective vaccine against the virus. mRNA-based vaccines have been extensively explored in the last few years for immune therapy applications and viral infections. Due to its negative charge and stability issues, mRNA molecules need suitable agents for intracellular delivery. LNPs are one of the most clinically advanced and commonly used tool for mRNA delivery [16].

In the present study, we developed LNPs-encapsulated mRNA as a vaccine platform. Endosomal escape of LNPs is a key step for endosomal release and functional activity of mRNA therapeutics. Towards this, several structurally-different lipids were chosen from our previous work. Lipid #2 and #5 contain hydrazine linker and lipids #10, #14 and #15 contains ethanolamine linker. However, the hydrophobic lipid tails chosen from fusogenic linoleic tails or acid sensitive mixed hydrophobic lipid tails. We compared these lipids in the form of LNPs for their ability to deliver mRNA and facilitate protein expression in mice, by employing commonly used routes of administration. First, we conducted a luciferase-based screen to evaluate protein expression efficiency and kinetics. IVIS data showed that two of these formulations, LNP #14 and #15, were more potent in terms of both level and duration of luciferase expression, as was measured by the luminescence signal after luciferin injection.

Next, we examined the immunologic response that developed against the expressed luciferase protein. The immunogenicity of the luciferase protein has been demonstrated previously [17], and we speculated that *in vivo* mRNA delivery efficiency would correlate with the resulting immunological responses. While the humoral response was rather limited, as one would expect after a single immunization, a substantial cellular response was recorded against the luciferase protein. Interestingly, immunization with LNP #2, led to a robust cellular immune response that was comparable to that of LNP #14 despite a lower and shorter-lived luminescent signal. This shows that one should take caution in making predictions regarding immunogenicity based on protein expression patterns. The elicitation of a significant immune response depends ultimately on the extent to which the delivered antigen is taken up by antigen presenting cells (APC), which migrate to lymph nodes and induce T cell activation and subsequent production of immune mediators. It is therefore possible, that in the case of LNP #2, while the IVIS data indicated a more limited pattern of luciferase expression, the tissues and cell types in which the protein was expressed, enabled the establishment of a more robust immune response. This issue will be further addressed in future studies that will evaluate tissue and organ-specific luciferase expression.

Given the robust cellular response observed in animals injected with LNPs #2 and #14, these formulations were used for the subsequent vaccination experiment with SARS-CoV-2 RBD mRNA. The RBD protein was chosen as an antigen for immunization based on recent data demonstrating the importance of the RBD domain in SARS-CoV-2 vaccine design, by elicitation of protective immunity by an RBD-based DNA vaccine [18] and a recombinant RBD-based vaccine [19].

Mice were vaccinated in a prime-boost regimen with naked RBD mRNA, LNP-encapsulated RBD mRNA or recombinant RBD.

Firstly, naked RBD mRNA immunization was unable to elicit detectable humoral or cellular responses before or after the boost, suggesting that the mRNA was most likely degraded and was incapable of triggering an effective immune response. Conversely, mice immunized with LNP RBD mRNA developed substantial anti-spike IgG titers, a robust cellular response, and high levels of neutralizing antibodies after boost administration in both intramuscular and intradermal groups. Interestingly, while the pre-boost humoral response was largely undetected in these immunization groups, ELISpot analysis demonstrated that a cellular response could be detected at that early stage, particularly in mice immunized intramuscularly, demonstrating the importance of characterization of the cellular response in SARS-CoV-2 immunization studies. While the two LNP formulations, #2 and #14, led to comparable humoral and cellular responses in intramuscularly-immunized mice, formulation #14 exhibited superior immunogenicity following intradermal administration of LNP RBD mRNA. This effect was most pronounced at the PRNT assay results (Figure 4C, middle panel). The observed cellular responses induced by LNP #14 may be attributed to their ability to activate APC via i.d. and i.m. administration, whereas LNP #2 is proficient via i.m. route only. This interesting observation suggests that not only LNPs structure but also route of administration needs to be considered for future clinical development of vaccines.

A large number of recombinant protein vaccines are currently in pre-clinical development, and several spike/RBD-based vaccines have entered clinical trials [2]. In order to evaluate the relative efficiency of our LNP-based mRNA vaccine, we performed a recombinant RBD immunization experiment in parallel, using a standard protein immunization protocol (a prime-boost s.c. administration of recombinant protein in the presence of an adjuvant) [20]. Although displaying comparable anti-RBD IgG and VSV neutralizing titers, the recombinant RBD immunization was unable to mount a significant cellular response as was recorded in the LNP RBD mRNA vaccine groups (Figure 4). These data demonstrate the inherent advantage of mRNA vaccination over recombinant protein vaccination in elicitation of immune response. While recombinant protein vaccination is dependent upon antigen uptake by APC, intracellular antigen expression following mRNA vaccination eventually leads to efficient peptide epitope MHC class I presentation which facilitates cytolytic T lymphocyte priming (in addition to helper T cell response). This combined activation of the two T cell subtypes yields a robust, long term humoral and cellular response, which may account for the apparent cellular response we observe following LNP RBD mRNA vaccination, and not after recombinant RBD immunization.

Two major concerns in the development of a safe SARS-CoV-2 vaccine is antibody-dependent enhancement (ADE) and vaccine-associated enhanced respiratory disease (VAERD), which could worsen the clinical manifestations of infection [21]. Since these two phenomena have been linked with a T helper 2 cell-biased response [21], we evaluated Th1 (IFNγ, IL-2) and Th2 (IL-4, IL-10) cytokine secretion in response to stimulation of splenocytes from vaccinated mice with SARS-CoV-2 spike. Our data demonstrate that both mRNA LNP formulations induced a Th1-biased cellular response towards the spike protein.

In summary, we report here an LNP-based RBD mRNA vaccine which was designed using a preliminary *in vivo* screen of structurally-different ionizable lipid-based LNPs. Several groups have recently reported evidence of immunogenicity and efficacy of LNP-based RBD mRNA vaccines [22–25], but have not referred to the issue of LNP lipid composition. The physicochemical properties of the ionizable lipids may have dramatic effects on delivery and protein expression efficiency, and should be considered upon LNP-based mRNA vaccine design.

## Materials and methods

### Animals

Female BALB/c mice (6–8 weeks old) were obtained from Charles River and randomly assigned into cages in groups of 10 animals. The mice were allowed free access to water and rodent diet (Harlan, Israel). All animal experiments were conducted in accordance with the guideline of the Israel Institute for Biological Research (IIBR) animal experiments committee. Protocol numbers: #M-60-19, #M-30-20.

### Production of SARS-CoV-2 antigens for immunization and *in vitro* assays

Recombinant SARS-CoV-2 spike glycoprotein, was expressed in pcDNA3.1^+^ plasmid, as recently described [26]. A stabilized soluble version of the spike protein (based on GenPept: QHD43416 ORF amino acids 1-1207) was designed to include proline substitutions at positions 986 and 987, and disruptive replacement of the furin cleavage site RRAR (residues at position 682-685) with GSAS. C-terminal his-tag as well as a strep-tag, were included in order to facilitate protein purification. Expression of the recombinant proteins was performed using ExpiCHO^TM^ Expression system (Thermoscientific, USA, Cat# A29133) following purification using HisTrap^TM^ (GE Healthcare, UK) and Strep-Tactin®XT (IBA, Germany). The purified protein was sterile-filtered and stored in PBS.

Human Fc-RBD fused protein was expressed using previously designed Fc-fused protein expression vector (Tal-Noy-Poral et al 2015), giving rise to a protein comprising of two RBD moieties (amino acids 331-524, see accession number of the S protein above) owing to the homodimeric human (gamma1) Fc domain (huFc). Expression of the recombinant proteins was performed using ExpiCHO^TM^ Expression system (Thermoscientific, USA) following purification using HiTrap Protein-A column (GE healthcare, UK). The purified protein was sterile-filtered and stored in PBS.

### mRNA

CleanCap® firefly luciferase mRNA was a kind gift from BioNtech RNA Pharmaceuticals (Mainz, Germany). CleanCap®, pseudouridine-substituted Fc-conjugated RBD mRNA (331-524 aa) was purchased from TriLink Bio Technologies (San Diego, CA, USA). The Fc-conjugated RBD mRNA was designed to include the exact translated Fc-RBD protein sequence as the recombinant protein.

### LNP preparation and characterization

LNPs were synthesized by mixing one volume of lipid mixture of ionizable lipid, DSPC, Cholesterol and DMG-PEG (40:10.5:47.5:2 mol ratio) in ethanol and three volumes of mRNA (1:16 w/w mRNA to lipid) in acetate buffer. Lipids and mRNA were injected in to a micro fluidic mixing device Nanoassemblr® (Precision Nanosystems, Vancouver BC) at a combined flow rate of 12 mL/min. The resultant mixture was dialyzed against phosphate buffered saline (PBS) (pH 7.4) for 16 h to remove ethanol.

Particles in PBS were analyzed for size and uniformity by dynamic light scattering (DLS). Zeta potential was determined using the Malvern zeta-sizer (Malvern Instruments Ltd., Worcestershire, UK). RNA encapsulation in LNPs was calculated according to Quant-iT RiboGreen RNA Assay Kit (Thermo Fisher, Waltham, MA, USA).

### Animal vaccination experiments

For *in vivo* LNP formulations screen, groups of 6-8-week-old female BALB/c mice were administered intramuscularly (50μl in both hind legs), intradermally (100μl) or subcutaneously (100μl) with luciferase mRNA (5μg) encapsulated with five different LNP formulations (LNPs #2, #5, #10, #14, #15). Luciferase expression was monitored as described in the bioluminescence imaging studies section. 28 days post-intramuscular injection, serum and spleen were collected from mice for evaluation of the immunologic response that developed towards luciferase.

For RBD mRNA vaccination studies, groups of 6-8-week-old female BALB/c mice were administered intramuscularly (50μl in both hind legs) or intradermally (100μl) with SARS-CoV-2 mRNA (5μg) encapsulated with LNP formulations #2 or #14.

For recombinant RBD (rRBD) vaccination studies, groups of 6-8-week-old female BALB/c mice were administered subcutaneously (100μl) with hFc-rRBD (10μg), hFc-rRBD emulsified in complete/incomplete Freund’s adjuvant (CFA/IFA), or adjuvant alone as control.

Both RBD mRNA-and recombinant RBD-immunized animals were boosted at day 25 with the same priming dose administered on day 0. Serum and spleens were collected on day 23 (“pre-boost”) and 49 (“post-boost”) for evaluation of immunologic response towards SARS-CoV-2 RBD, and measurement of cytokine secretion.

### Bioluminescence Imaging Studies

Bioluminescence imaging was performed with an IVIS Spectrum imaging system (Caliper Life Sciences). Female BALB/c mice were administered D-luciferin (Regis Technologies) at a dose of 150 mg/kg intraperitoneally. Mice were anesthetized after receiving D-luciferin with a mixture of ketamine (60 mg/kg) and xylazine (10 mg/kg) and placed on the imaging platform. Mice were imaged at 5 minutes post administration of D-luciferin using an exposure time of 60 seconds. Bioluminescence values were quantified by measuring photon flux (photons/second) in the region of interest using the Living IMAGE Software provided by Caliper.

### ELISA

ELISA was performed for the detection of luciferase-or SARS-CoV-2 spike-specific antibodies in immunized mouse sera. MaxiSORP ELISA plates (Nunc) were pre-coated with recombinant luciferase (0.4μg/ml, Promega, #E1701) or spike protein (2μg/ml) overnight at 4°C in carbonate buffer. Plated were washed three times with PBST (PBS+0.05% Tween-20) and blocked with 2% BSA (Sigma-Aldrich, #A8022) in PBST for 1 h at 37 °C. After three washes with PBST, plates were incubated with serially diluted mouse sera for 1 h at 37 °C. Following washing, goat anti-mouse alkaline phosphatase-conjugated IgG (Jackson Immuno Research Labs, #115-055-003) was added for 1 h at 37 °C. The plates were washed with PBST and reactions were developed with p-nitrophenyl phosphate substrate (PNPP, Sigma-Aldrich, N2765). Plates were read at 405 nm absorbance and antibody titers were calculated as the highest serum dilution with an OD value above 2 times the average OD of the negative controls.

### Cytokine assays

Splenocytes from immunized mice were incubated in the presence of SARS-CoV-2 spike protein (10μg/ml). Culture supernatants were harvested 48 h later and analyzed for cytokines by ELISA techniques with commercially available kits. IL-2 (DY402), IL-4 (DY404) and IL-10 (DY417) kits were obtained from R & D Systems, Minneapolis, Minn.

### Murine IFNγ ELISpot Assay

Mice spleens were dissociated in GentleMACS C-tubes (Miltenyi Biotec), filtered, treated with Red Blood Cell Lysing Buffer (Sigma-Aldrich, #R7757), and washed. Pellets were resuspended in 1ml of CTL-Test™ Medium (*CTL, #CTLT 005*) supplemented with 1% fresh glutamine, and 1 mM Pen/Strep (Biological Industries, Israel), and single cell suspensions were seeded into 96-well, high-protein-binding, PVDF filter plates at 400,000 cells/well. Mice were tested individually in duplicates by stimulation with recombinant luciferase (13μg/ml, Promega, #E1701), SARS-CoV-2 spike protein (10μg/ml), Concanavalin A (Sigma-Aldrich, #0412) (2μg/ml) as positive control, or CTL medium as negative control (no antigen). Cells were incubated with antigens for 24 h, and the frequency of IFNγ-secreting cells was determined using Murine IFNγ Single-Color Enzymatic ELISPOT kit (CTL, #MIFNG 1M/5) with strict adherence to the manufacturer’s instructions. Spot forming units (SFU) were counted using an automated ELISpot counter (Cellular Technology Limited).

### Plaque Reduction Neutralization Test (PRNT)

VSV-spike ^[27]^stocks were prepared by infection of Vero E6 cells for several days. When viral cytopathic effect (CPE) was observed, media were collected, clarified by centrifugation, aliquoted and stored at −80°C. Titer of stock was determined by plaque assay using Vero E6 cells.

For plaque reduction neutralization test (PRNT), Vero E6 cells (0.5*10^6^ cells/well in 12-well plates) were cultured in DMEM supplemented with 10% FCS, MEM non-essential amino acids, 2nM L-Glutamine, 100 Units/ml Penicillin, 0.1 mg/ml streptomycin and 12.5 Units/ml Nystatin (Biological Industries, Israel) overnight at 37°C, 5% CO_2_.

Serum samples were 3-fold serially diluted (ranging from 1:50-1:12,500) in 400 μl of MEM supplemented with 2% FCS, MEM non-essential amino acids, 2nM L-Glutamine, 100 Units/ml Penicillin, 0.1 mg/ml streptomycin and 12.5 Units/ml Nystatin. 400 μl containing 300 PFU/ml of VSV-spike were then added to the diluted serum samples and the mixture was incubated at 37 °C, 5% CO_2_ for 1 h. Monolayers were then washed once with DMEM w/o FBS and 200 μl of each serum-virus mixture was added in triplicates to the cells for 1 h. Virus mixture without serum was used as control. 1 ml overlay [MEM containing 2% FBS and 0.4% tragacanth (Sigma, Israel)] was added to each well and plates were incubated at 37 °C, 5%CO_2_ for 72 h. The number of plaques in each well was determined following media aspiration, cells fixation and staining with 1 ml of crystal violet (Biological Industries, Israel). NT50 was defined as serum dilution at which the plaque number was reduced by 50%, compared to plaque number of the control (in the absence of serum).

### Transmission Electron Microscopy Analysis

A drop of aqueous solution containing LNPs was placed on the carbon-coated copper grid and dried. The morphology of LNPs was analyzed by a JEOL 1200 EX (Japan) transmission electron microscope

### Statistical analysis

All values are presented as mean plus standard error of the mean (s.e.m). Antibody titers, neutralizing titers, ELISpot data and cytokine levels were compared using two-way ANOVAs or t-tests as depicted in the figure captions. All statistical analyses were performed using GraphPad Prism 8 statistical software.

## Acknowledgments

This work was supported by the Lewis Trust grant to D.P.

## Data availability

All relevant data are available from the authors upon reasonable request.

