## Supplemental Materials and Methods for "Design of SARS-CoV-2 RBD mRNA Vaccine Using Novel Ionizable Lipids"

**Supplementary information**

Linoleic alcohol obtained from TCI chemicals and all other chemicals obtained from sigma-Aldrich unless otherwise mentioned. Lipid **2**, **5**, **10** and **14** were synthesized according to the procedure mentioned in our previous work [12]. The lipid **15** synthesized according to the procedure described below.

### Synthesis of Lipid 15:

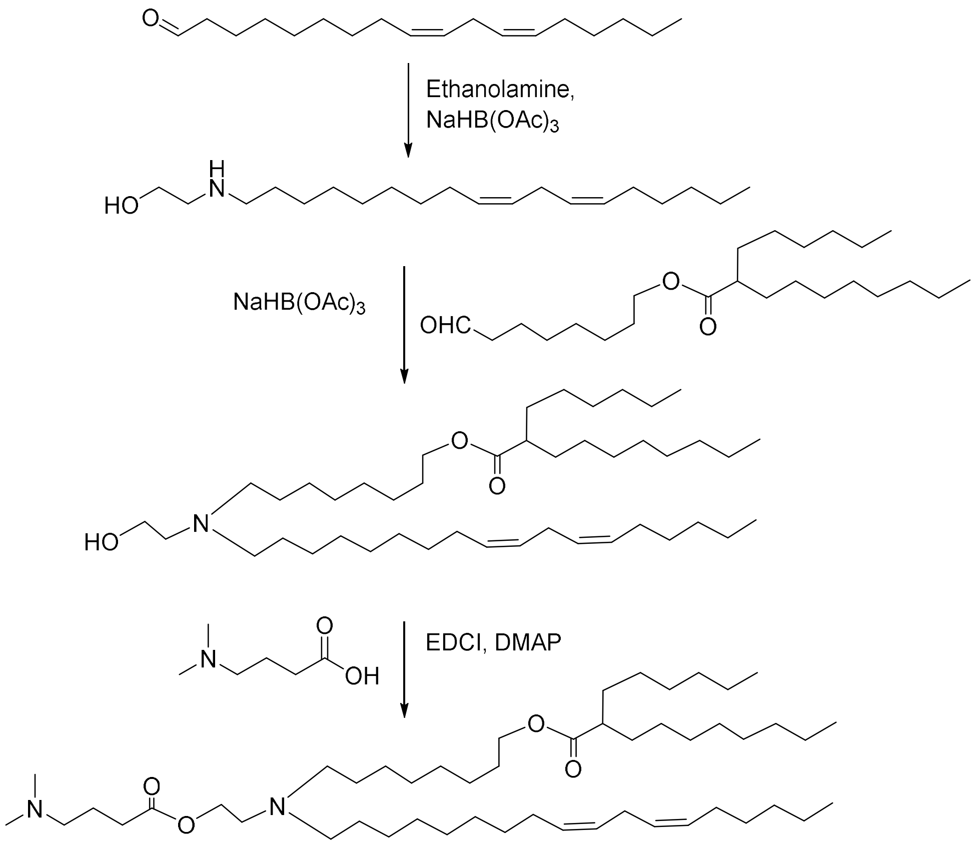


**Lipid 15**

***2-(((9Z, 12Z)-Octadeca-9, 12-dien-1-yl) amino) ethan-1-ol***

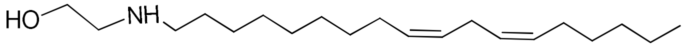

Chemical Formula: $C_{20}H_{39}NO$
Molecular Weight: 309.5298

Linoleyl aldehyde synthesized previously [12] (3.2g, 12.12 mmol) and ethanolamine (0.89ml, 14.54 mmol) were dissolved in dry $CH_2Cl_2$ (60 mL) under nitrogen atmosphere and stirred for 1hr at room temperature. Then sodium triacetoxyborohydride (5.1g, 24.24 mmol) was added and stirred for 16hr at room temperature. The reaction mixture was quenched with sat.$NaHCO_3$ solution followed by extracted with DCM. The organic layer was washed with brine solution and dried over anhydrous $Na_2SO_4$. The solvent was evaporated and the residue was purified by column chromatography using 0-10% MeOH in $CHCl_3$ to get desired compound (2.0 g, 54 %) as a pale yellowish liquid.

$^1$H NMR (400 MHz, $CDCl_3$): $\delta$ 5.40-5.30 (4 H, m), 3.64 (2 H, t, $J$ = 5.2 Hz), 2.83-2.73 (4 H, m), 2.62 (2 H, t, $J$ = 7.2 Hz), 2.04 (4 H, q, $J$ = 6.8 Hz), 1.54-1.43 (2 H, m), 1.41-1.21 (16 H, m), 0.88 (3 H, t, $J$ = 6.8 HZ).

ESI-MS: *m/z* 310.5 $[M+1]^+$

***8-((2-Hydroxyethyl)((9Z,12Z)-octadeca-9,12-dien-1-yl)amino)octyl 2-hexyldecanoate***

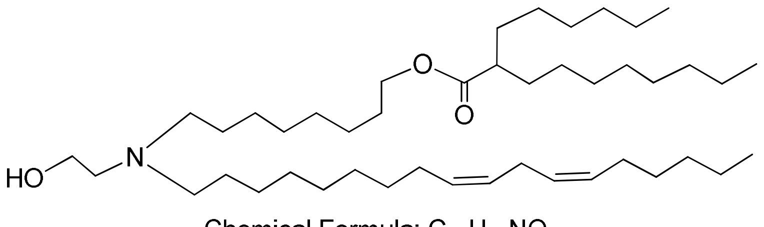

Chemical Formula: $C_{44}H_{85}NO_3$
Molecular Weight: 676.1506

The above linoleyl ethanolamine (411mg, 1.33mmol) and previously synthesized 8-oxooctyl 2-hexyldecanoate [12] (61 mg, 1.60 mmol) were dissolved in dry $CH_2Cl_2$ (30 mL) under nitrogen atmosphere and stirred for 1hr at room temperature. Then sodium triacetoxyborohydride (56 mg, 2.66 mmol) was added to the reaction mixture and stirred for 24hr. The reaction mixture was quenched with sat.$NaHCO_3$ solution then extracted with DCM. The organic layer was washed with brine solution and dried over anhydrous $Na_2SO_4$. The solvent was evaporated and the residue was purified by column chromatography using 0-2% MeOH in $CHCl_3$ yielded 760 mg (86%) of pure lipid as pale yellowish oil.

$^1$H NMR (400 MHz, $CDCl_3$): $\delta$ 5.43-5.27 (4 H, m), 4.06 (2 H, t, $J$ = 6.4 Hz), 3.55 (2 H, t, $J$ = 5.2 Hz), 3.08 (1 H, br), 2.77 (2 H, t, $J$ = 6.4 Hz), 2.61(2 H, t, $J$ = 5.2 Hz), 2.48 (4 H, t, $J$ = 7.6 Hz), 2.35-2.25 (1 H, m), 2.04 (4 H, q, $J$ = 6.8 Hz), 1.67-1.51 (4 H, m), 1.50-1.38 (6 H, m), 1.38-1.16 (44 H, m), 0.88 (3 H, t, $J$ = 6.8 HZ), 0.87 (6 H, t, $J$ = 6.8 HZ).

ESI-MS: *m/z* 676.9 $[M+1]^+$

***8-((2-((4-(Dimethylamino)butanoyl)oxy)ethyl)((9Z,12Z)-octadeca-9,12-dien-1-yl)amino)octyl 2-hexyldecanoate (LIPID 15)***

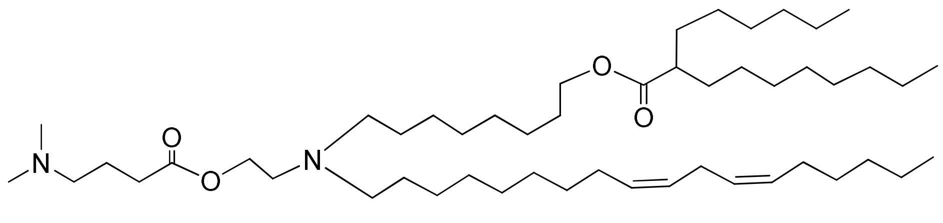

**EA-502**

Chemical Formula: $C_{50}H_{96}N_2O_4$

Molecular Weight: 789.3082

The above alcohol (750mg, 1.11mmol), *N, N*-dimethyl aminobutyric acid hydrochloride (371mg, 2.22mmol), EDC.HCl (424mg, 2.22 mmol) and DMAP (13mg, 0.11mmol) were dissolved in dry $CH_2Cl_2$ (30 mL) under argon atmosphere and stirred for 24hr at room temperature. Excess DCM was added and the reaction mixture washed with brine solution followed by dried over anhydrous $Na_2SO_4$. The organic solvent was evaporated and the residue was purified by column chromatography using 0-5% Isopropanol in $CHCl_3$ to get pure **Lipid 15** (665mg, 76%) as pale yellow color oil.

$^1$H NMR (400 MHz, $CDCl_3$): $\delta$ 5.43-5.27 (4 H, m), 4.11 (2 H, t, $J$ = 6.4 Hz), 4.05 (2 H, t, $J$ = 6.8 Hz), 2.77 (2 H, t, $J$ = 6.4 Hz), 2.67 (2 H, t, $J$ = 5.2 Hz), 2.43 (4 H, t, $J$ = 7.6 Hz), 2.37-2.26 (5 H, m), 2.22 (6 H, s), 2.04 (4 H, q, $J$ = 6.8 Hz), 1.78 (2 H, quint, $J$ = 7.2 Hz), 1.66-1.51 (4 H, m), 1.48-1.16 (50 H, m), 0.88 (3 H, t, $J$ = 6.8 Hz), 0.87 (6 H, t, $J$ = 6.8 Hz).

ESI-MS: *m/z* 790.1 $[M+1]^+$; 395.6 $[M/2+1]^+$

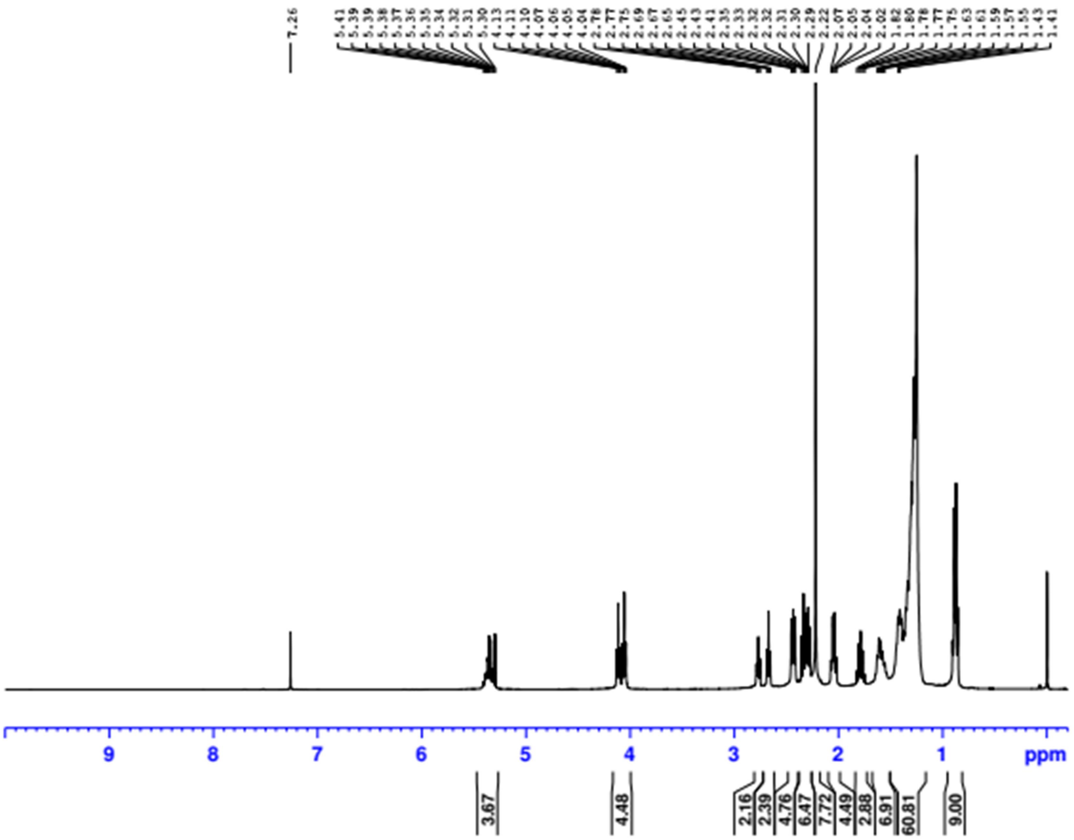
7.26
5.41 5.39 5.39 5.38 5.37 5.36 5.35 5.34 5.32 5.31 5.30 4.13 4.11 4.10 4.07 4.06 4.05 4.04 2.78 2.77 2.75 2.69 2.67 2.65 2.45 2.43 2.41 2.35 2.33 2.32 2.32 2.31 2.30 2.29 2.22 2.07 2.05 2.04 2.02 1.82 1.80 1.78 1.77 1.75 1.63 1.61 1.59 1.57 1.55 1.43 1.41
9 8 7 6 5 4 3 2 1 ppm
3.67
4.48
2.16 2.39 4.76 6.47 7.72 4.49 2.88 6.91 60.81
9.00

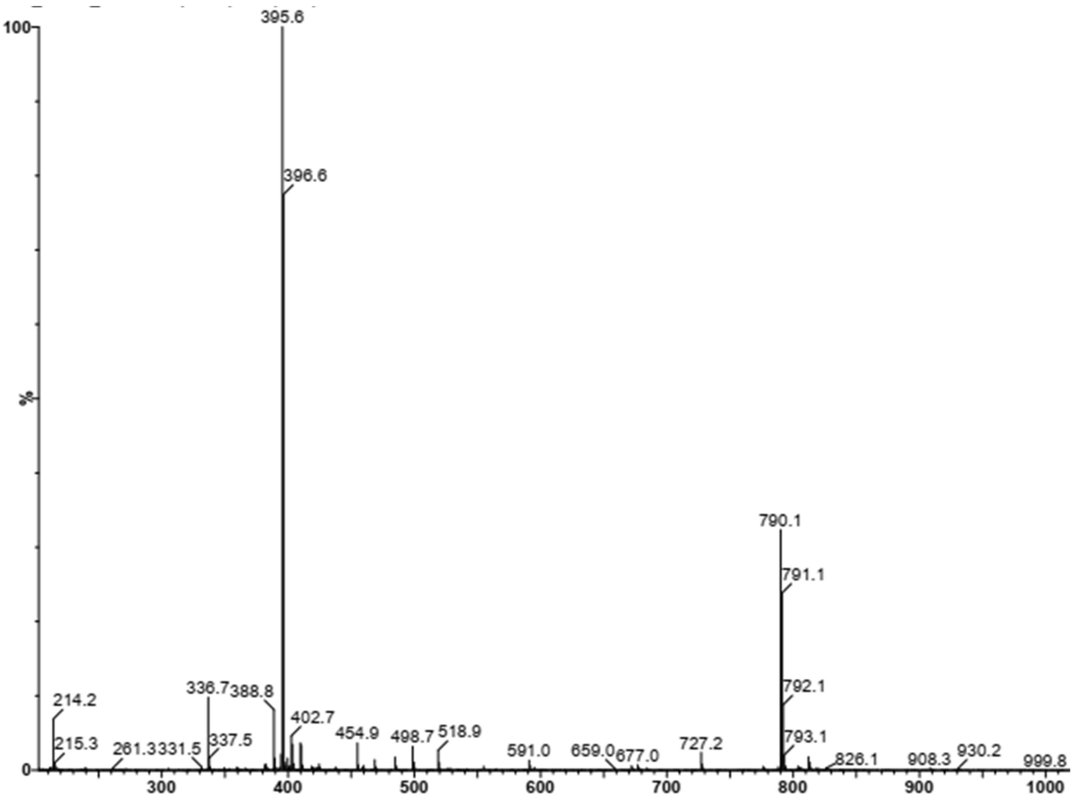
100
%
0
395.6
396.6
790.1
791.1
792.1
793.1
214.2
215.3
261.3
331.5
336.7
337.5
388.8
402.7
454.9
498.7
518.9
591.0
659.0
677.0
727.2
826.1
908.3
930.2
999.8
300 400 500 600 700 800 900 1000
